# Patterns of Neural Activation During an Initial Social Stress Encounter are Predictive of Future Susceptibility or Resilience: A FosTRAP2 Study

**DOI:** 10.1101/2023.03.10.532130

**Authors:** Dalia Murra, Kathryn L. Hilde, Huzefa Khalil, Stanley J. Watson, Huda Akil

## Abstract

Repeated social stress is a significant factor in triggering depression in vulnerable individuals, and genetic and environmental factors interact to contribute to this vulnerability. Interestingly, the role of experience in shaping vulnerability is not well studied. To what extent does an individual’s initial reaction to a given stressor influence their response to similar stressors in the future? And how is this initial response encoded at the neural level to bias towards future susceptibility or resilience?

The Chronic Social Defeat Stress (CSDS) mouse model offers an ideal opportunity to address these questions. Following 10 days of repeated social defeat, mice diverge into two distinct populations of social reactivity: resilient (interactive) and susceptible (avoidant). It is notable that the CSDS paradigm traditionally uses genetically inbred mice, indicating that this divergence is not genetically determined. Furthermore, the emergence of the two phenotypes only occurs following several days of exposure to stress, suggesting that the repeated experience of social defeat influences future susceptibility or resilience.

In this study, we asked whether specific patterns of neural activation during the initial exposure to the social defeat stress can predict whether an individual will eventually emerge as resilient or susceptible. To address this question, we used Fos-TRAP2 mouse technology to capture brain-wide neural activation patterns elicited during the initial stress exposure, while allowing the mice to go on to experience the full course of CSDS and diverge into resilient and susceptible populations. Using a high-throughput brain-wide cell counting approach, we identified the bed nucleus of the stria terminalis and lateral septal nucleus as key hubs for encoding social defeat. We also identified the basomedial amygdala as a hub for encoding future susceptibility, and the hippocampal CA1 area and medial habenula for encoding future resilience.

Our findings demonstrate that the initial experience with social stress induces a distinct brain-wide pattern of neural activation associated with defeat, as well as unique activation patterns that appear to set the stage for future resilience or susceptibility. This highly orchestrated response to defeat is seen especially in animals that emerge as resilient compared to susceptible. Overall, our work represents a critical starting place for elucidating mechanisms whereby early experiences can shape vulnerability to affective disorders.

## INTRODUCTION

Many social animals, including humans, rely on social interactions as a key source of survival. Navigating social interactions is a highly complex process that involves paying attention to the environmental context, perceiving cues from social partners, and responding with appropriate emotional control. When negative social encounters are a constant source of stress in the environment, the propensity to develop vulnerability increases (Hofmann 2007; Kocovski et al. 2005). However, the perception of social stress is subjective, and there is substantial variety in outcomes of stress across individuals. Prior social experiences may play a crucial role in an individual’s perception of and response to future social events. While the cumulative effects of chronic social stress on the brain and body have been well-studied, less is known about how the initial exposure can shape the final affective state. Early patterns of responsiveness, such as coping strategies and activation of brain-wide circuits, likely set the stage for future social reactivity (Palamarchuk and Vaillancourt 2021; Sinha et al. 2016; Snyder et al. 2015). Here we ask whether there are predictive neural activation patterns present early during the stress response that act as a risk factor in the outcome of stress reactivity.

Any given stressful encounter can provoke a range of neural and behavioral responses, and these initial responses may in turn shape future outcomes of subsequent stressors(Evans and Kim 2013; Folkman and Moskowitz 2004; McEwen 1998). In this way, the distinctive neural and behavioral activity pattern caused by the initial stress event may predispose the development of either susceptibility or resilience to future stress. In mice, the chronic social defeat stress paradigm (CSDS) is a widely validated model of individual variation in response to social stress (Hammels et al. 2015; Krishnan et al. 2007). In this paradigm, inbred C57BL6/J intruder mice are repeatedly subjected to bouts of physical and psychosocial stress by a larger, aggressive resident CD1 mouse. Following CSDS, the intruder mice diverge into two distinct populations of social reactivity: a socially interactive “Resilient” group and a socially avoidant “Susceptible” group (Golden et al. 2011). While it has been established that several days of social defeat are necessary before the Resilient and Susceptible classification emerge (Bagot et al. 2015; Kudryavtseva 1994; Wells et al. 2017), work from our lab found that behavioral coping strategies displayed during the initial stress encounter predict whether mice develop resilient or susceptible responses in later social interaction testing (Murra et al. 2022). The unique behavioral responses individuals exhibit during their first experience of stress are likely to reflect distinct patterns of underlying neural activity that then set the stage for future resilience or susceptibility.

Several brain regions known to play a role in major depressive disorder (MDD) have also been implicated in mediating reactivity to social stress, including the prelimbic cortex (PrL), nucleus accumbens (NAc), amygdala (AMY), ventral hippocampus (vHipp), and ventral tegmental area (VTA) (Mayberg et al. 2005; Nestler et al. 2002). However, whether this neural circuitry differs prior to exposure to chronic social defeat, or whether it emerges as a result of the repeated stress, remains unclear. Two key studies have addressed this question in the context of the CSDS model. One study relied on *in vivo* fiber photometry and found that a baseline increase in D1-MSN activity in the NAc was a predictive marker of future resilience but not susceptibility (Muir et al. 2018). Another study relied on *in vivo* local field potential recordings in the MDD-network brain regions and found that before stress, subtle differences in oscillatory signatures mapped onto future resilience versus susceptibility (Hultman et al. 2018). Other circuit-based studies have focused on assessing neural activity after the stress has occurred (Anacker et al. 2016; Bagot et al. 2015; Chaudhury et al. 2013; Heshmati et al. 2020; Hultman et al. 2018; Muir et al. 2020; Nam et al. 2019; Vialou et al. 2010). Each of these previous studies have selected individual brain regions *a priori* or assessed specific microcircuits in their contributions to social outcome. The practical challenges therefore reside in 1) scalability of assessing unbiased brain-wide neural response patterns and 2) the fact that classification as Susceptible or Resilient requires a multi-day course of daily social stress.

While classic immediate early gene studies have been useful in mapping neuronal activation patterns post-mortem (add refs for classic fos studies from CSDS), recent advances in genetic circuit tracing enables *in vivo* capture of neuronal activity. Indeed, using the Fos-TRAP system, we can now capture brain-wide neural activity patterns in living animals during the first episode of social defeat and allow them to live on to undergo the chronic stress experience and be further classified into Resilient or Susceptible groups. The FosTRAP2 mouse line links a promotor-driven Fos Cre-ERT2 system in a tamoxifen-dependent manner to an LSL-tdTomato reporter to allow for permanent brain-wide labeling of activated neurons restricted to a specific time window (DeNardo et al. 2019; Guenthner et al. 2013) (**Fig. 1A**). Using this system, we labeled and quantified brain-wide ensembles of neurons that are activated on Day 1 of stress to permanently ‘TRAP’ the stress-naïve circuit. To that end, our overall hypothesis was that brain-wide activation patterns that emerge during the initial defeat encounter can predict the eventual split in social reactivity. Using a high-throughput brain-wide cell counting approach, we identified the bed nucleus of the stria terminalis and lateral septal nucleus as key hubs for encoding social defeat. We also identified the basomedial amygdala as a hub for encoding future susceptibility, and the hippocampal CA1 area and medial habenula for encoding future resilience. Overall, we found evidence for selective brain regions and networks that are activated on Day 1 that are specific to Defeat alone, and others that distinguish between groups that will eventually emerge as Resilient and Susceptible.

**Figure 1.**
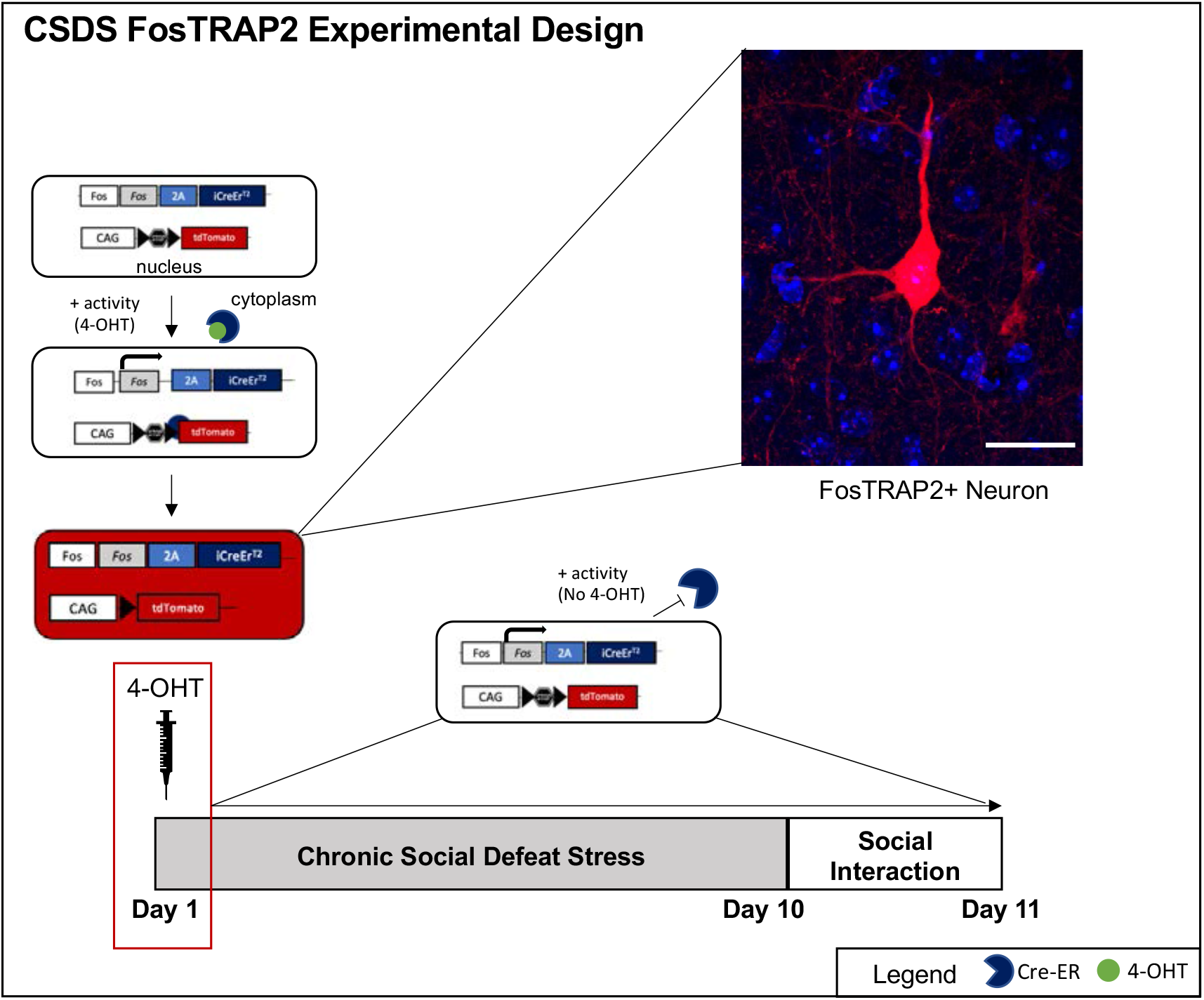
FosTRAP2 CSDS Approach. The TRAP system requires two transgenes: one that expresses CreERT2 from an activity-dependent Fos promoter and one that allows expression of tdTomato, in a Cre-dependent manner. To capture neural circuitry during the first day of stress, we injected 4-hydroxytamoxifen (4-OHT) 1 hour following 5 minutes of defeat. When 4-OHT is present, Cre-ER recombination can occur in active cells, allowing for an indelible tomato tag. See the visualization of a FosTRAP2+ neuron (imaged at 40X). Red= FosTRAP+, blue=DAPI. Scale bar = 20μm. Following Day 1, no 4-OHT is present, and no recombination occurs. For the remainder of the paradigm, theCReERt2 is retained in the cytoplasm of active cells in which its expressed.

## METHODS

### Animals

Transgenic FosTRAP2 (Jackson Laboratory; Fos2mt.1(iCreERT2)Luo/J; strain #(030323)), and Ai14D(Jackson Laboratory; B6l;129S6-Gt(ROSA)26sortm14(CAG-tdtomato)Hze/J; Strain #007908) mouse lines were crossed in-house to generate FosTRAP2-Ai14 mice (*hereafter referred to as FosTRAP2*). In total, 32 FosTRAP2 male mice (10-12 weeks of age) were used for the Social Defeat and were housed 2-4 per cage until start of experiment. 36 male CD1 retired breeders (3-6 months) from Charles River Laboratories (Wilmington, MA) were used as aggressors and social interaction stimulus mice. For both CD1s and FosTRAP2 mice, enrichment in the form of enviro-paks was provided. Housing rooms were kept under a 12 h light: 12 h dark cycle starting at 7 AM. Mice were provided with mouse chow and tap water *ab libitum*, in accordance with the *NIH Guidelines for the Care and Use of Laboratory Animals. All animal protocols utilized were approved by the University of Michigan Institute of Animal Use Care and Use Committee (IAUCAC)*.

### Chronic social defeat stress (CSDS) and Control Rotation (CR)

CSDS and CR experiments were conducted for 10 days using a previously described protocol (Golden et al. 2011). All testing occurred between 9-11am each day. Prior to use in the CSDS paradigm, CD1 mice were screened for their latency to attack (<1 min. interval to attack for 2 consecutive days). During CSDS, FosTRAP2 mice were subjected to 5-minute bouts of agonistic encounters from a novel, aggressive CD1 mouse. After each session, FosTRAP2 mice were placed across a perforated plexiglass divider from the CD1 in the resident CD1’s cage (27×48×15cm). During CR, FosTRAP2 mice were placed across from one another in the same cage set up as CSDS but had no physical interaction. CR mice were rotated across to a new partner each day. FosTRAP2 mice were singly housed and placed in static cages following CSDS or CR.

### Social Interaction Test (SI)

Following ten days of CSDS, subjects were tested for SI (morning of 11^th^ day). The SI test took place in an open field arena (41×41×40cm), which had a removable wire mesh stimulus cage (10×10×10cm) against one of the sidewalls. The interaction zone was defined as a 14×24cm rectangle that surrounded the cage. SI was assessed with two sequential trials, each recorded and tracked using automatic tracking software (Noldus Ethovision XT; Noldus Technology, Leesburg, VA). The first trial consisted of the FosTRAP2 mouse exploring the novel arena with an empty cage for 2.5 minutes. For the second trial, a novel, aggressive CD1 mouse was placed in the cage, and the FosTRAP2 mouse was returned to the arena for another 2.5 minutes. The SI ratio was calculated as the time spent in the interaction zone with the CD1 present divided by the time spent in the interaction zone with the CD1 absent. We categorized the mice based on their SI ratio: < 1 indicated susceptibility and ≥ 1 indicated resilience.

### Tamoxifen

1 hour following the 5-minute Defeat encounter on Day 1, FosTRAP2 mice were injected with 4-hydroxy-tamoxifen (4-OHT) intraperitoneal at a dose of 50mg/kg. 4TM (Sigma H6278) was dissolved in an aqueous solution containing 10% DMSO and 10% Tween-80 in saline. All cages were changed the following day to avoid the re-uptake of 4-hydroxy-tamoxifen.

### Histology

FosTRAP2-Ai14 mice were perfused transcardially one day following social interaction (day 12) with cold 1X PBS solution followed by 4% PFA. Brains were post-fixed in 4% PFA overnight and dehydrated with 20% sucrose, then flash-frozen in -20°C isopentane. Coronal sections were collected in a series of 10 at 20µM using a freezing cryostat at -24°C and immediately mounted to the glass slide. Slides were stored at -80°C until processing. The sampling rate was every 200µM starting anterior at bregma 1.98 and ending posterior at bregma -3.08. Slides were washed 3×5-min with 1X PBS and then incubated with DAPI 1:10,000 for 10 min. Slides were then washed 3×5min, and coverslipped using Invitrogen™ Prolong™ Gold Antifade Mountant (Cat#P10144).

### Imaging Parameters

All images were taken on an Olympus IX83 Inverted Microscope at a 10x objective and imaged at 546nm (FosTRAP2+ cells) and 405nm (DAPI).

### Cell Counting and Registration

Individual cell bodies for FosTRAP2+ cells were identified using the automated cell detection feature in NeuroInfo (MBF Bioscience, Williston, VT). For these cells, the largest cell diameter detected was 40µM, and the smallest was 6µM at an intensity of 16-26 nearest to the background (depending on the section). Our dependent variable in all analyses were the number of FosTRAP2+ cells. The NeuroInfo ‘Registration’ feature was used on the DAPI channel to register individual sections to the Allen Mouse Brain Atlas.

### Statistical Analysis Software Use

Graphpad Prism and R (version 4.0.2) were used for figure design and statistical analyses. In R, *pastecs, psych, nlme, pheatmap, and emmeans* packages were used for visual and statistical analyses. *Cor* function was used to assess correlations between all brain regions within Control, Defeat, Resilient, and Susceptible groups. *Pheatmap* was used to visualize clusters of correlations between brain regions and groups. Raw cell counts were log2 transformed prior to assessing group differences. The *nlme* package with the *lm* function was used to run linear regressions on all brain regions between groups and post-hoc Tukey multiple comparisons was done using the *emmeans* package. All significance thresholds were set at *p≤0*.*05* and FDR at *<*0.1.

### Detailed Analysis Pipeline

Our analysis pipeline is split into three separate phases: A. Data Input, B. Group Comparisons, and C. Network Analysis.

#### Data Input

To assess differences between the initial Control vs. Defeat, as well as future Resilient vs. Susceptible populations, we used two different data input approaches (**Fig. 2A**). Our first level of analysis used an unbiased exploratory approach: We selected 73 brain regions based on two criteria (see **Table S1)**: 1) include any region that had FosTRAP2 activity (cell count of >1 across all animals in at least one group) during the stress or social experience on Day 1t and 2) exclude any region that was specific to sensory or visual processing. This exclusion criteria was based on the fact that when 4-OHT reaches the brain, it can persist for up to 6 hours (Guenthner et al. 2013). Therefore, while the injection was timed for peak Tomato+ labeling following stress, the activity patterns that are captured likely include both the agonistic encounter itself, as well as the psychosocial stress and sensory feedback that can persist during the first day of either Defeat or social encounter in the Control condition. Our next level of analysis utilized a candidate approach: We selected 26 brain regions (See **Table S2)** based on circuitry that has been previously identified in the CSDS model, as well as areas known in the literature to be involved in stress and social responses. These 26 brain regions were a subset of the 73 regions used in our first analysis approach.

**Figure 2.**
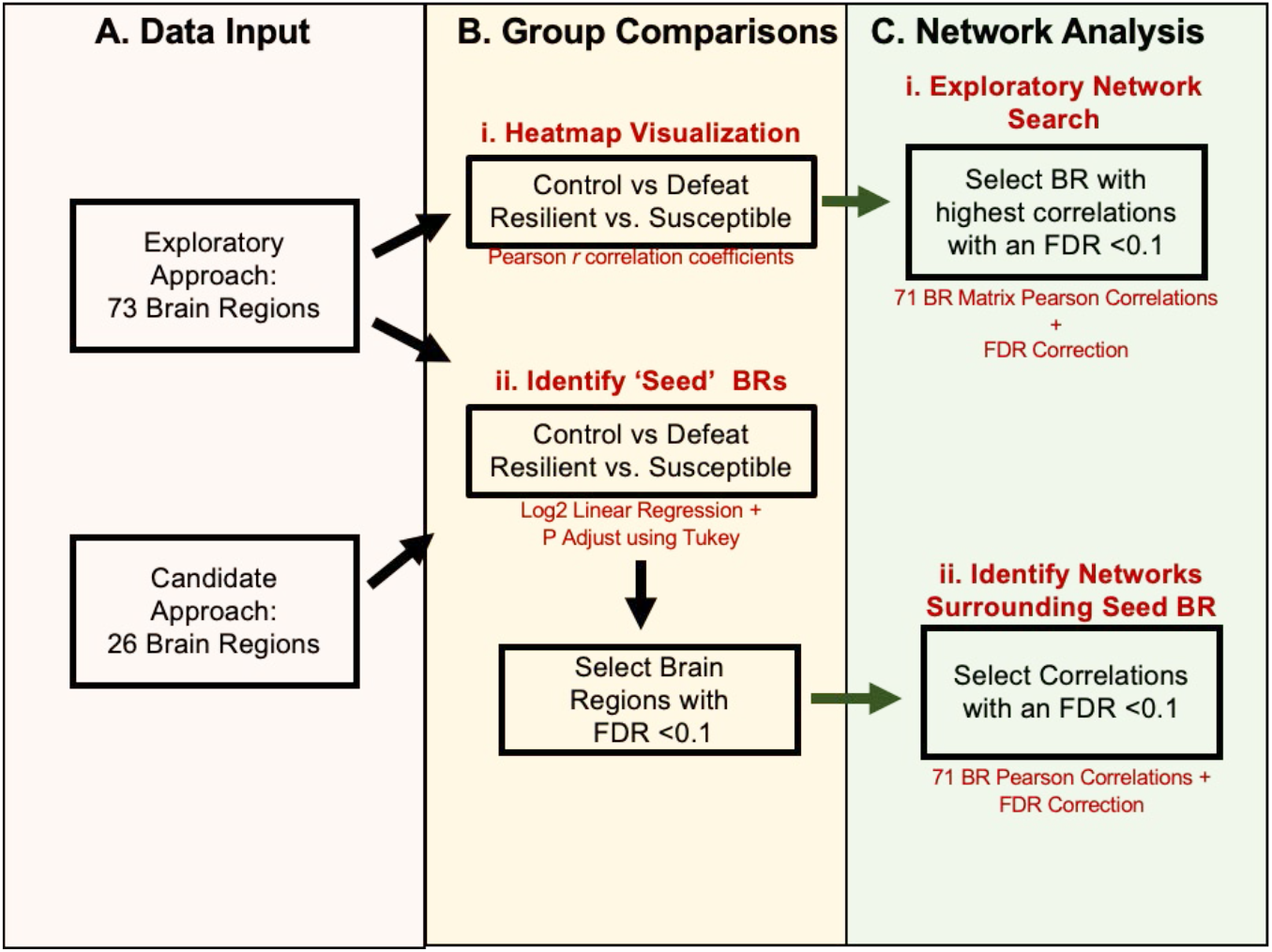
Data Analysis Pipeline. Data Analysis was split into 3 sections A. Data Input, B. Group Comparisons, C. Network Analysis

#### Group Comparisons

<u>We compared brain-wide activity patterns between groups with both descriptive and inferential statistical approaches (Fig 2B). For a descriptive comparison of activity patterns, we used</u> the Pearson *r* correlations in activation levels between all 73 brain regions and generated heatmaps for each group (**Fig 2B.i.)**. To determine if FosTRAP2+ activity levels were significantly different between groups, we first log2 transformed the FosTRAP2+ cell counts and then ran a linear regression (*lm* function in *nlme* R package) for each brain region, correcting for multiple comparisons using the Tukey-HSD test (*emmeans* function and package in R). These direct group comparisons were done for both exploratory (73 brain regions) and candidate (26 brain regions) approaches (**Fig 2B.ii)**.

#### Network Analysis

Our network analyses also used both an exploratory and candidate approach. In our exploratory network approach, we created a matrix of Pearson *r* correlations of all 73 brain regions within each group and ran a Benjamini-Hoshberg correction on the dataset to correct for multiple comparisons. As such, we selected this exploratory seed brain region based on highest number of brain regions that correlated with the seed and met an FDR cut-off of <0.1 (**3B.i**.). In our candidate approach, we identified networks that surrounded the seed brain regions identified from our group comparisons in **2B.ii. (3B.ii.)**. Similar to the exploratory approach, we selected the ‘seed’ brain regions and ran a Pearson correlation analysis between the seed region and the other 72 surrounding brain regions. From there, we selected the brain regions that were most highly correlated with each seed region based on an FDR of <0.1.

## RESULTS

### Characterizing the Behavioral and Neural Response to CSDS

In order to assess neural circuitry associated with the initial Defeat encounter, we injected FosTRAP2 mice with 4-OHT on Day 1 of Social Defeat. FosTRAP2 mice experienced the traditional trajectory of all 10 days of social Defeat and were then placed in the social interaction test for final classification **(Fig. 1)**. Control mice demonstrated a preference for social interaction, with a mean SI Ratio of 1.311 (*sd=*+/-0.292). In contrast, mice that experienced social Defeat displayed a significant reduction in SI Ratio, with a mean of 0.688 (*sd*=+/-0.430) (**Fig. 3**; *p*=0.001, Mann-Whitney U=27). Within the Social Defeat group, 6 animals were defined as Resilient (socially interactive).

**Figure 3.**
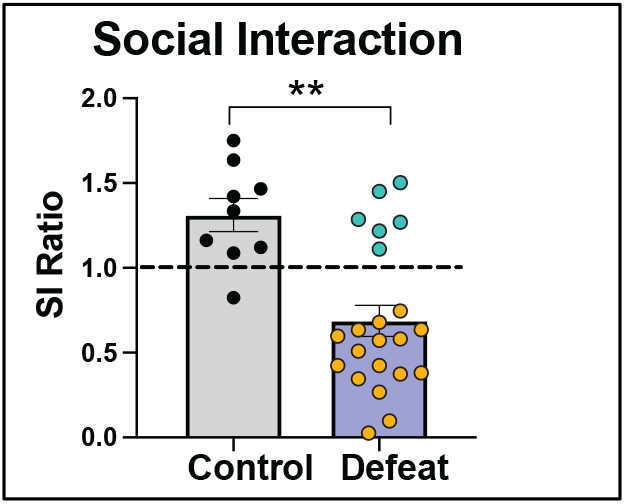
Chronic Social Defeat Stress in FosTRAP2 Mice. Social Interaction (SI): Defeated mice show a significant reduction in SI compared to controls (p=0.001, Mann-Whitney U=27).

We first wanted to assess whether there were differences in networks of activation between the initial Control (social) and Defeat (social stress) conditions. To do so, we took the Pearson *r* correlations between all brain regions to visualize overall differences in the network between groups. On day 1 of Defeat or Control rotation, defeated mice exhibited a more coordinated network of brain region activations, as indicated in the heatmaps by the increased positive Pearson *r* values (more red) (**Fig. 4A, B)**.

**Figure 4.**
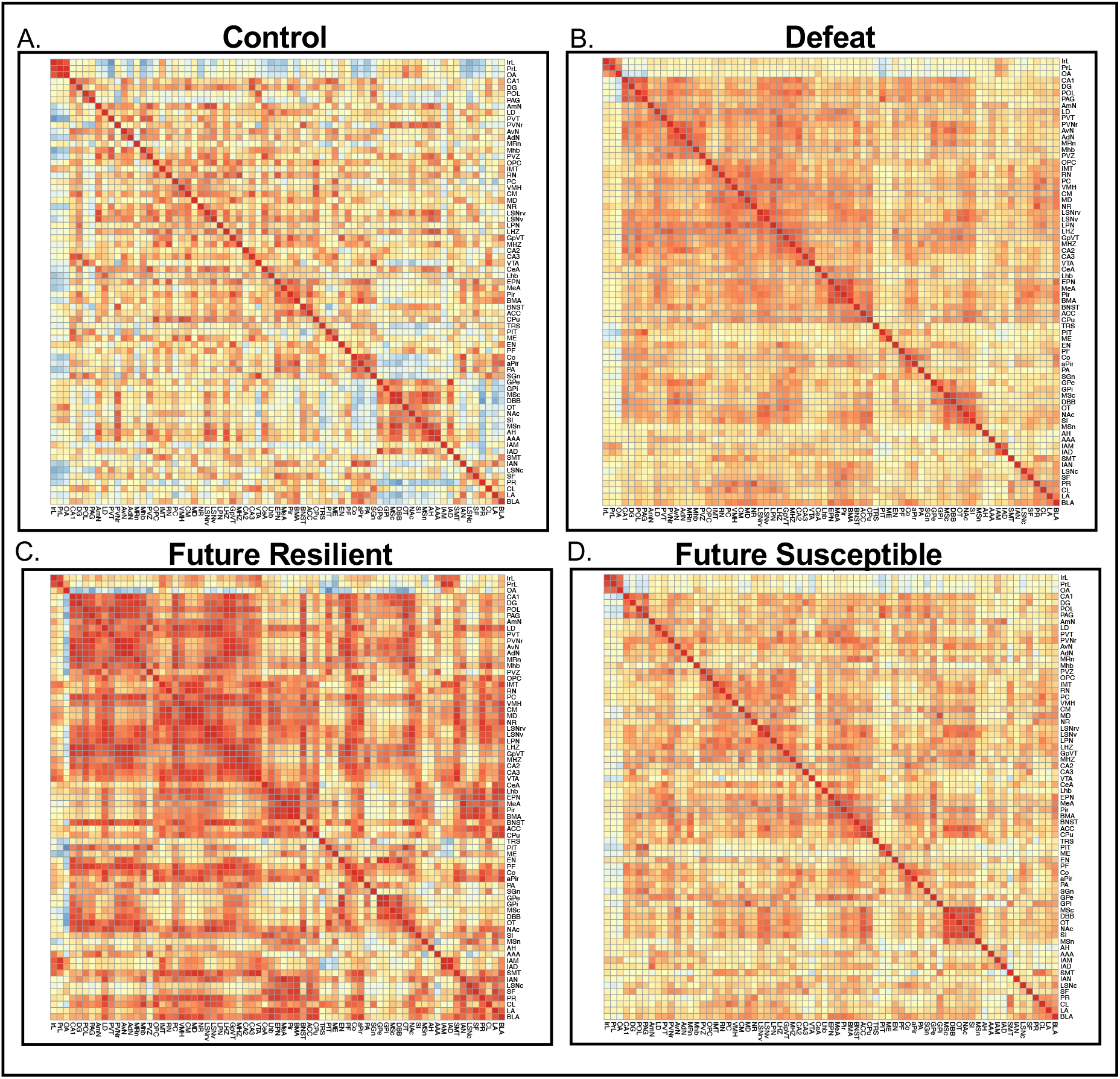
Heatmap Visualization of Correlations within A. Control, B. All Defeated, C. Future Resilient, and D. Future Susceptible mice. Colors in heatmap indicate a range of Pearson *r* correlations, more red=1, more blue=-1. Order of brain regions was based on a clustering of all groups combined. All Pearson *r* correlations are based on total number of FosTRAP+ cells within each group for each brain region. Each row represents a brain region and its Pearson correlation with other brain regions.

Within the Defeated group, future Resilient and Susceptible mice appeared to have different network patterns of activation. Indeed, visually comparing the correlation heatmaps generated for the two groups, Resilient mice had more positive correlations between brain regions in comparison to Susceptible (increased red, positive Pearson *r* values) (**Fig. 4C, D)**. This difference was not due to the differences in number of animals in each group, as a random selection of n=5 from each group condition resulted in a similar visual pattern (**Fig. S1)**. The initial Defeat encounter resulted in an increased positive correlation network of activation compared to controls, which was possibly primarily driven by the increase of correlations within the Resilient group. Thus, *resilient mice exhibit a more coordinated network of activation than susceptible mice*.

### An Exploratory Network Analysis

Our first network-level approach was to rank order the FDR corrected p-values of the correlation matrices within each group to determine which brain region emerges as the most significantly correlated. This allowed us to gain an unbiased view to determine which brain region could be deemed as a “seed” region of interest within each group.

#### The Lateral Septum is a Seed Region in Control and Defeat Conditions

Within the Control group, the lateral septal nucleus, rostroventral part (LSNrv) was the most significantly correlated with other areas (21 brain regions). Similarly, within the defeated group, the lateral septal nucleus, ventral part (LSNv) was also the most significantly correlated region (will be expanded upon in **Fig. 7**). This finding indicates that the lateral septum (LS) may play a unique role in Day 1 stress and social behaviors, possibly mediated by distinct sub-regions of the LS. In addition, this further emphasizes our identification of the LSNv as a key region that differentiates the defeat experience.

#### The Basomedial Amygdala and Hippocampal CA1 Seed Networks are Potential Candidates in Predicting Future Social Outcome

Importantly, this analysis revealed that future Resilient and Susceptible mice exhibited distinct seeds. In future Resilient mice, the hippocampal CA1 (CA1) region correlated significantly with 22 brain regions (**Fig 5A)** In future Susceptible mice, the basomedial amygdala (BMA) correlated significantly with 52 brain regions **Fig. 5B**). This suggests that there distinct seed regions underly the different patterns of correlations associated with Resilient and Susceptible outcomes.

**Figure 5.**
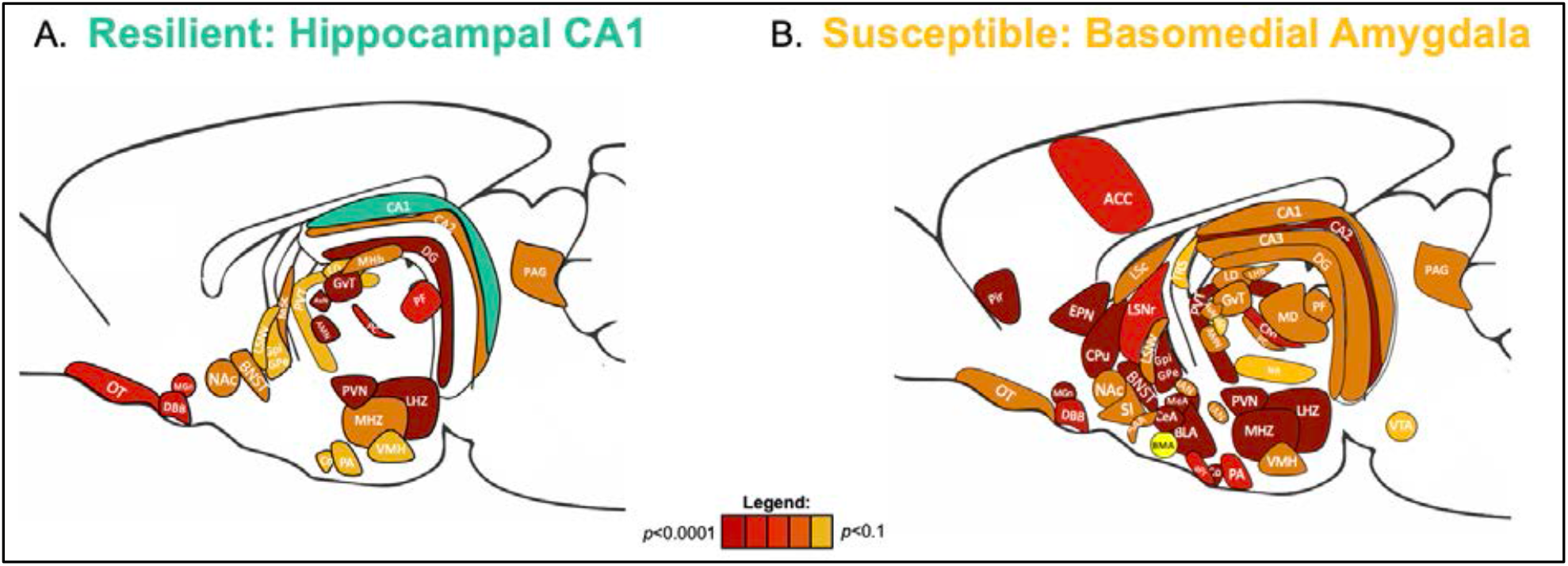
CA1 and BMA Future Resilient and Susceptible Exploratory Seed Network Maps. A. Brain regions correlating with the future Resilient CA1 seed (seed is in blue-green). B. Brain regions correlating with the future Susceptible BMA seed (seed is in bright yellow). These networks reflect a range of significantly correlated brain regions with seeds, from p <0.001 (red) to p<0.1 (yellow).

### A Targeted Group Comparison Approach

#### Control vs. Defeat: The LSNv and Bed Nucleus of the Stria Terminalis are the Most Differentially Activated Brain Regions

When comparing Defeated vs. Control mice, 6/73 regions examined showed statistically significant differences in neural activation (See **Table 1**). Four of these brain regions met an FDR cut-off of below 0.1: An increase in the LSNv (*p*=0.004, *FDR*=0.025), bed nucleus of the stria terminalis (BNST) (*p*=0.0007, FDR=0.025), Medial Hypothalamic Zone (*p*= 0.006, FDR =0.092), and anterior hypothalamic nucleus (AH) (*p*=0.006, FDR=0.0918) was seen in Defeat compared to Controls. These differences are depicted in **Table 1** and **Figures 6C and D**. Similarly, in our candidate approach comprising of 26 brain regions, the top two significantly different regions between Control and Defeated mice that met an FDR cut-off of 0.05 were also the LSNv (*p=*0.000, *FDR*=0.0122) and BNST (*p=*0.000, *FDR*=0.017) (see **Table 2** and **Figures 6C and D)**. Our results indicate that activation of the BNST and LSNv distinguished animals that have experienced defeat compared to the control rotation.

**Table 1.** Top 10 Brain Regions from Exploratory Approach

| Region | Estimate | SE | df | t.ratio | P.value | FDR |
| --- | --- | --- | --- | --- | --- | --- |
| LSNv | 0.9423 | 0.2369 | 29 | 3.9781 | 0.0004 | 0.0249 |
| BNST | 0.6861 | 0.1804 | 29 | 3.8028 | 0.0007 | 0.0249 |
| MHZ | 0.5021 | 0.1693 | 29 | 2.9652 | 0.0060 | 0.0918 |
| AH | 0.6421 | 0.2179 | 29 | 2.9461 | 0.0063 | 0.0918 |
| MHb | 1.0836 | 0.3820 | 29 | 2.8368 | 0.0082 | 0.1001 |
| PVN | 0.5607 | 0.2093 | 29 | 2.6788 | 0.0120 | 0.1256 |
| EPN | 0.4073 | 0.2008 | 29 | 2.0282 | 0.0518 | 0.4486 |
| CA1 | 0.6090 | 0.3050 | 29 | 1.9968 | 0.0553 | 0.4486 |
| MeA | 0.3296 | 0.1719 | 29 | 1.9172 | 0.0651 | 0.4753 |

Red meets FDR cut-off, Brown is nominally significant, and Black is non-significant.

**Table 2.**
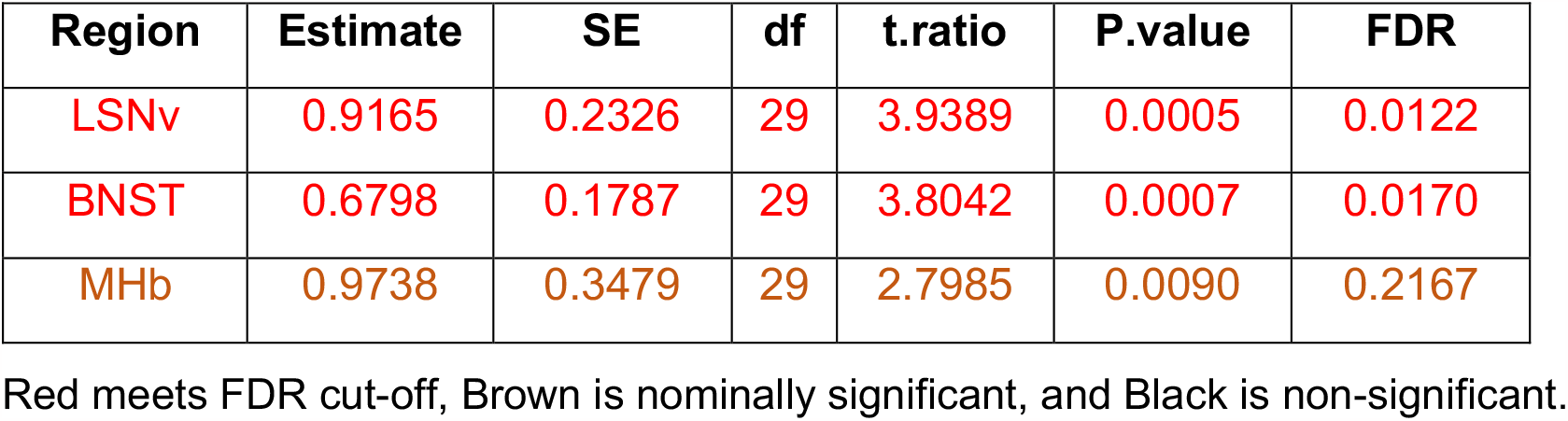
Top 3 Brain Regions from Candidate Approach in Control vs. Defeat

| Region | Estimate | SE | df | t.ratio | P.value | FDR |
| --- | --- | --- | --- | --- | --- | --- |
| LSNv | 0.9165 | 0.2326 | 29 | 3.9389 | 0.0005 | 0.0122 |
| BNST | 0.6798 | 0.1787 | 29 | 3.8042 | 0.0007 | 0.0170 |
| MHb | 0.9738 | 0.3479 | 29 | 2.7985 | 0.0090 | 0.2167 |

Red meets FDR cut-off, Brown is nominally significant, and Black is non-significant.

**Figure 6.**
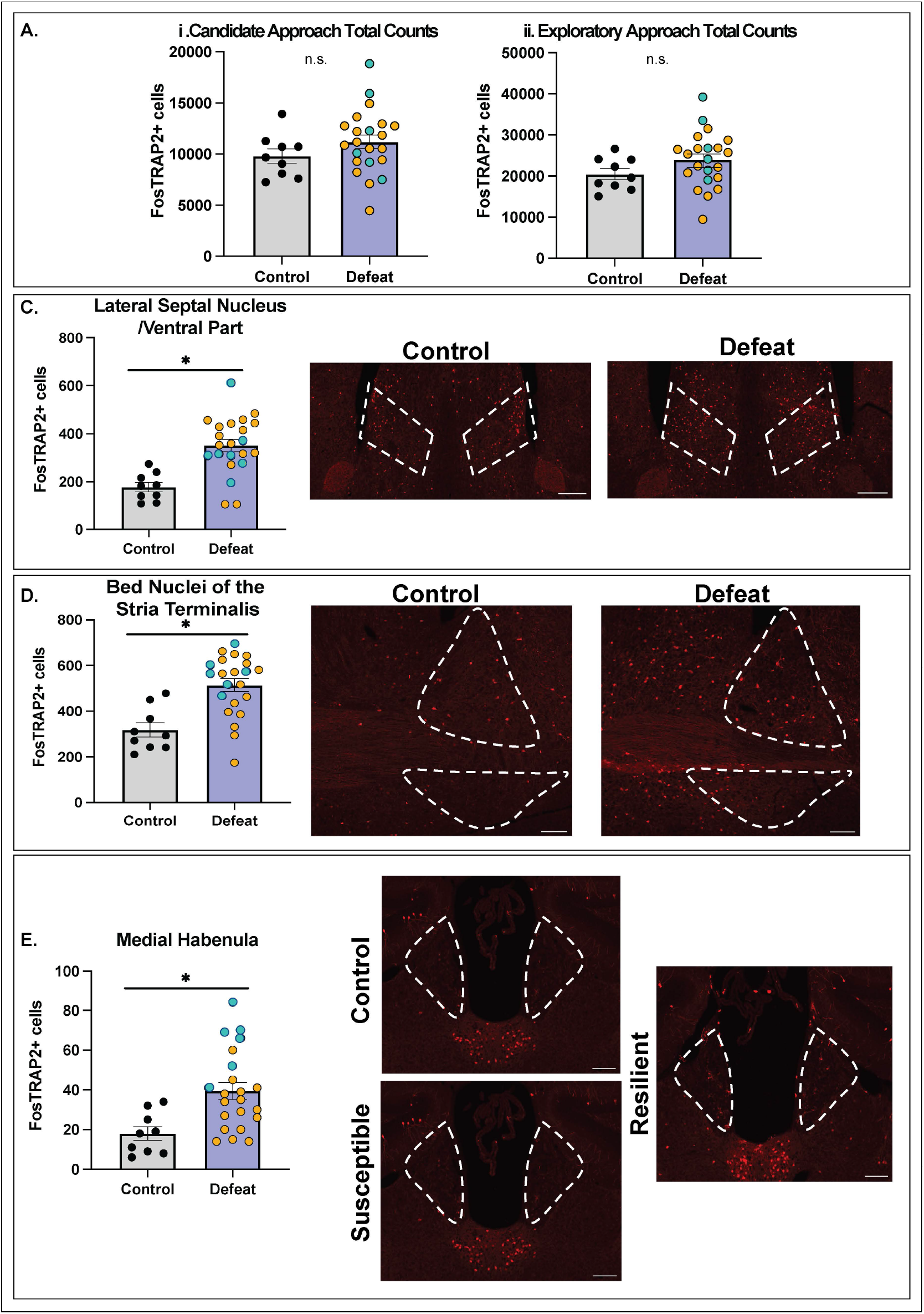
Brain Regions Selectively Active on Day 1. A. Total counts from the exploratory (73 brain regions) or candidate (26 brain regions) approach between Defeat and Control, and Future Resilient and Susceptible groups. There were no significant differences between Control vs. Defeat or Control vs. Resilient or Control vs. Susceptible or Susceptible vs. Resilient (p>0.05). B. Defeated animals had an increase of FosTRAP+ compared to controls (imaged at 4X). C. Defeated animals had an increase of FosTRAP+ cells in the BNST compared to controls (imaged at 10X). D. Resilient animals had an increase of FosTRAP+ cells compared to control and susceptible animals (Imaged at 10X). All scale bars are 100μm and all images were taken in stacks and Z-projected. Brain Regions of interest are outlined with white dashed line.

**Figure 7.**
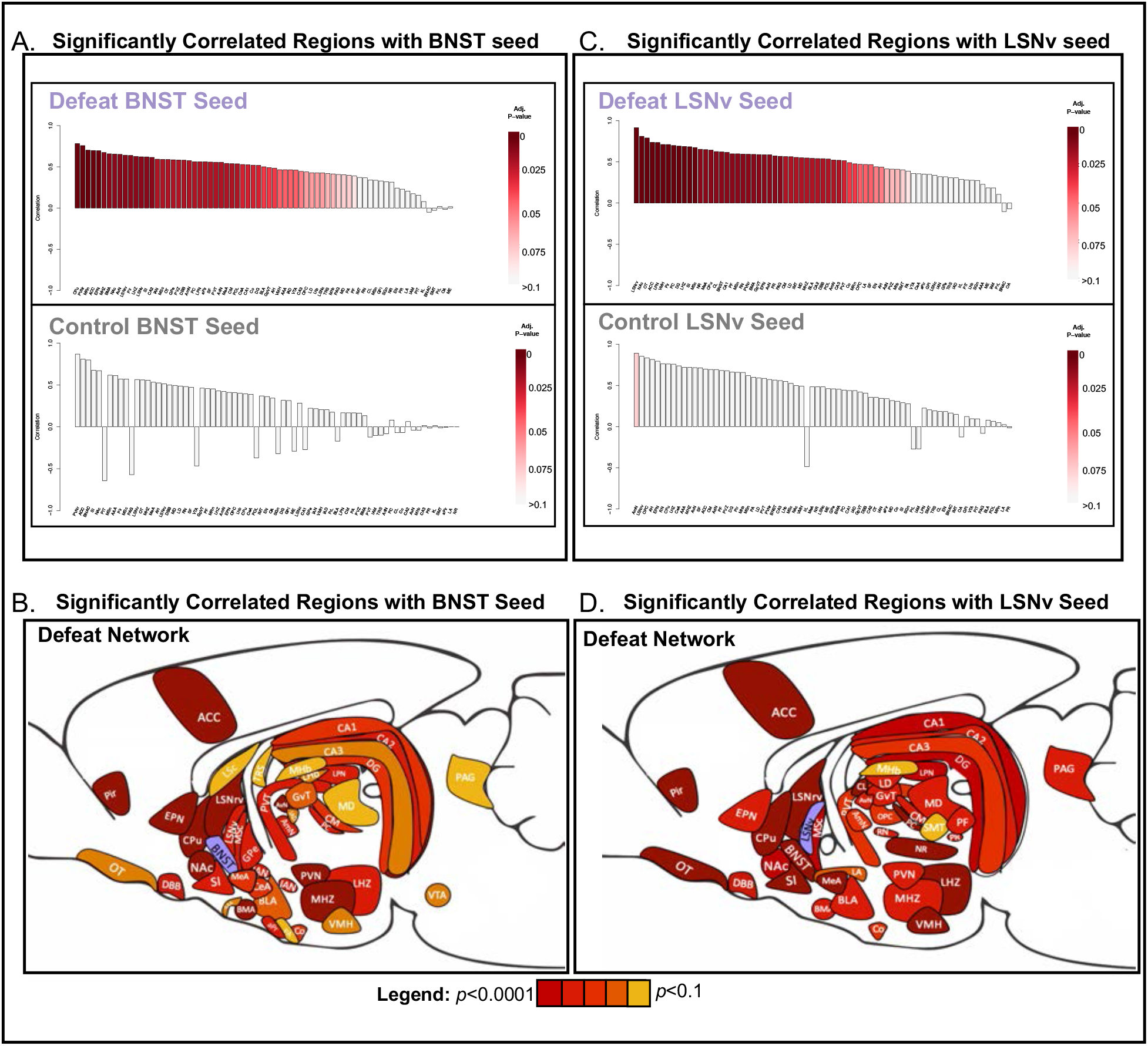
Significantly Correlated Brain Regions with LSNv and BNST Seed Networks. A. BNST Pearson R correlations of brain regions ranked based on adjusted p-values. B. Correlation network visualization of the BNST seed with all other significantly correlated regions in Defeated mice C. LSNv Pearson R correlations of brain regions ranked based on adjusted p-values D. Correlation network visualization of the LSNv seed with all other significantly correlated regions in Defeated mice. Color Code: Purple represents the defeat seed, and brain regions range from dark red (p<0.0001) to yellow (p<0.1) based on FDR significance.

#### Future Resilient vs. Susceptible: The Medial Habenula is the Most Differentially Activated Region

We next asked whether there were brain regions that were differentially activated in susceptible versus resilient animals. Four of 73 regions were nominally significant, with the medial habenula (MHb) showing the largest elevation in FosTRAP2+ cells in Resilient compared to Susceptible mice (**Table 3)**. Using the candidate approach, the medial habenula was also significantly different between groups, with Resilient mice showing greater activation than Susceptible mice (**Table 3.4, Fig. 6E;** *p*=0.003, *FDR*=0.0731). Our results indicate that as early as the first Defeat encounter, activation patterns within the MHb distinguish future Resilient and Susceptible mice.

**Table 3.**
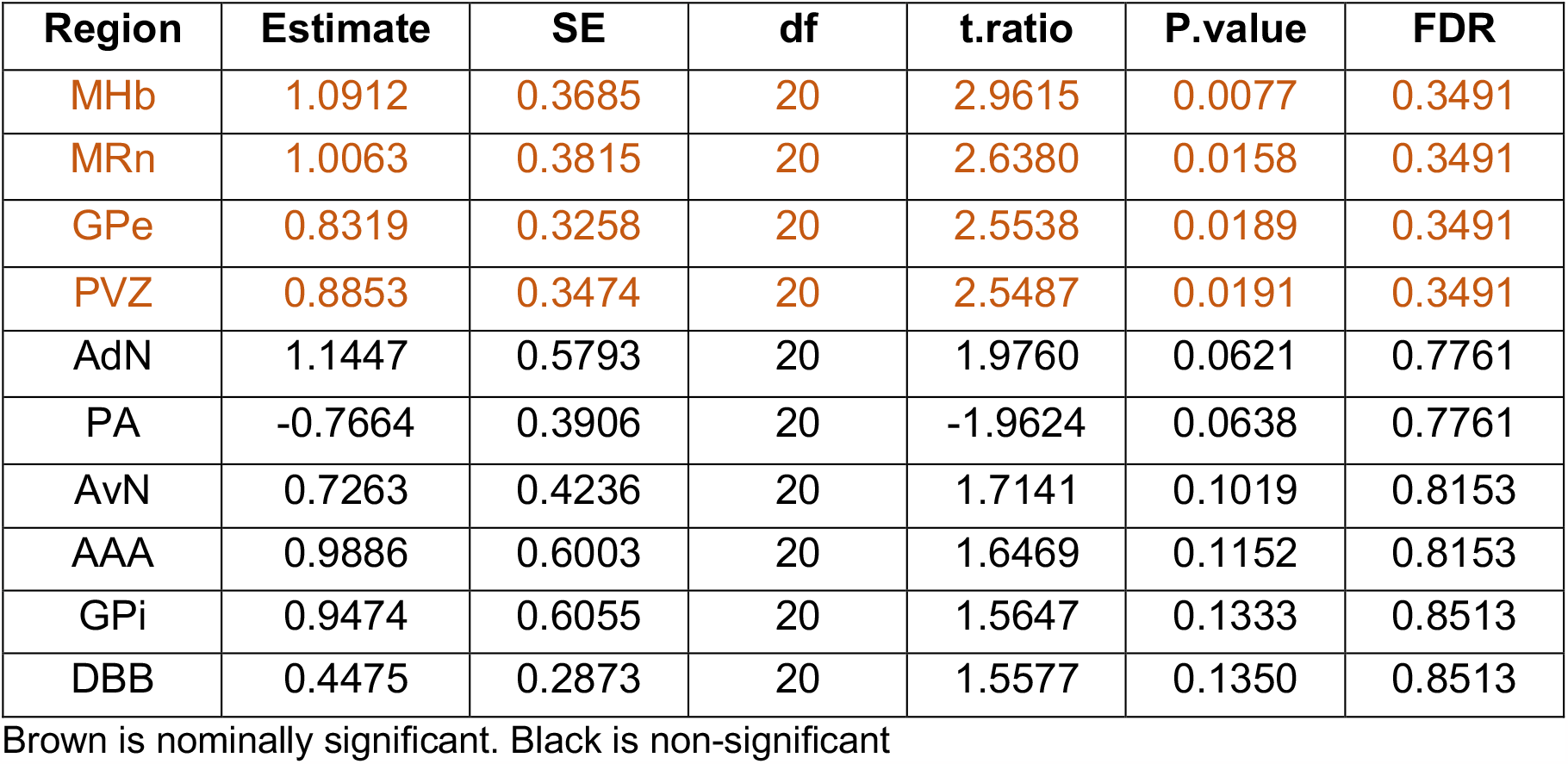
Top 10 Brain Regions from Exploratory Approach in Future Resilient and Susceptible Mice

**Table 4.** Top 3 Brain Regions from Candidate Approach in Future Resilient Mice

| Region | Estimate | SE | df | t.ratio | P.value | FDR |
| --- | --- | --- | --- | --- | --- | --- |
| MHb | 1.0887 | 0.2966 | 28 | 3.6703 | 0.00281 | 0.0731 |
| PVZ | 0.8806 | 0.3225 | 28 | 2.7302 | 0.02828 | 0.7071 |
| CeA | 0.4861 | 0.3033 | 28 | 1.6028 | 0.2611 | 1 |

Red meets FDR cut-off, Brown is nominally significant. Black is non-significant

### Network Analyses Using the BNST, LSNv, and MHb as Seeds

While we identified brain regions that were selectively activated by Defeat or future Resilient populations, we next wanted to ask whether there were other brain regions that potentially activated in unison to reveal circuit-specific differences. As these regions were the most differentially activated between groups, in the Control and Defeat conditions, we selected the LSNv and BNST as ‘seeds’ and in Resilient and Susceptible conditions, selected the MHb as a ‘seed’. We then correlated these seed regions with the other 72 brain regions and selected for networks of activation if the brain region Pearson *r* correlation met an FDR cut-off of <0.1.

#### The BNST and LSNv Reveal Significantly Correlated Networks of Activation in Defeated mice

The BNST in Control and Defeated mice correlated with different brain regions (**Fig. 7A**). Within Defeated mice, 52 brain regions met an FDR cut-off of 0.1 and significantly correlated with the BNST (**Fig. 7A,B** and see **Table S3** for statistics). In contrast, in Control mice, there were no correlations between the BNST and other brain regions that met an FDR cut-off of 0.1 (although 4 brain regions were nominally significant, see **Table S4** for statistics). This is consistent with the fact that BNST activity was selectively enhanced by Defeat, and suggests that the BNST was a centerpiece in an entire network of activation triggered by the Defeat stress. Most notably, in the Day 1 Defeat condition, the hippocampus, thalamic, hypothalamic, striatal, septal, olfactory, and cortical areas were significantly correlated with BNST activity (**Figure 6B** and see **Table S3** for full list of brain regions and statistics). Overall, compared to controls, defeated mice engage an additional network of activation that is highly integrated with the BNST.

Similarly, the LSNv in Control and Defeated mice had different patterns of correlations to other brain regions **(Fig. 7C)**. Within Defeated mice, 53 brain regions met an FDR cut-off of 0.1 and significantly correlated with the LSNv (**Fig. 6C, D** and see **Table S5** for statistics). In contrast, in Control mice, only the LSNv and other brain regions met an FDR cut-off of 0.1 (**Fig. 7C** and see **Table S6)**. Here again, the Defeat group showed a selective increase in LSNv activity compared to Controls, and that region is nodal to a network of activation that is unique to that group. In the Day 1 Defeat condition, the hippocampus, thalamic, hypothalamic, striatal, septal, olfactory, and anterior cingulate cortex were significantly correlated with LSNv activity (**Fig. 7D** and **Table S5**). Therefore, defeated mice have engaged a highly coordinated LSNv network.

#### The Medial Habenula (MHb) Differentiates Future Resilient Mice

The MHb in future Susceptible and Resilient mice had different strengths and levels of significance across brain regions (**Fig. 8A)**. Within Resilient mice, 4 brain regions met an FDR cut-off of 0.1 and significantly correlated with the MHb (**Fig. 7B;** see **Table S7** for statistics). In contrast, in Control mice, there were no correlations between the MHb and other brain regions that met an FDR cut-off of 0.1 (though 8 brain regions were nominally significant, see **Fig. 8A** and see **Table S8** for statistics). Overall, the MHb showed a selective increase in activity in the future Resilient group compared to Susceptible, and this increase led to a unique network of activation that is specific to future resilience to stress.

**Figure 8.**
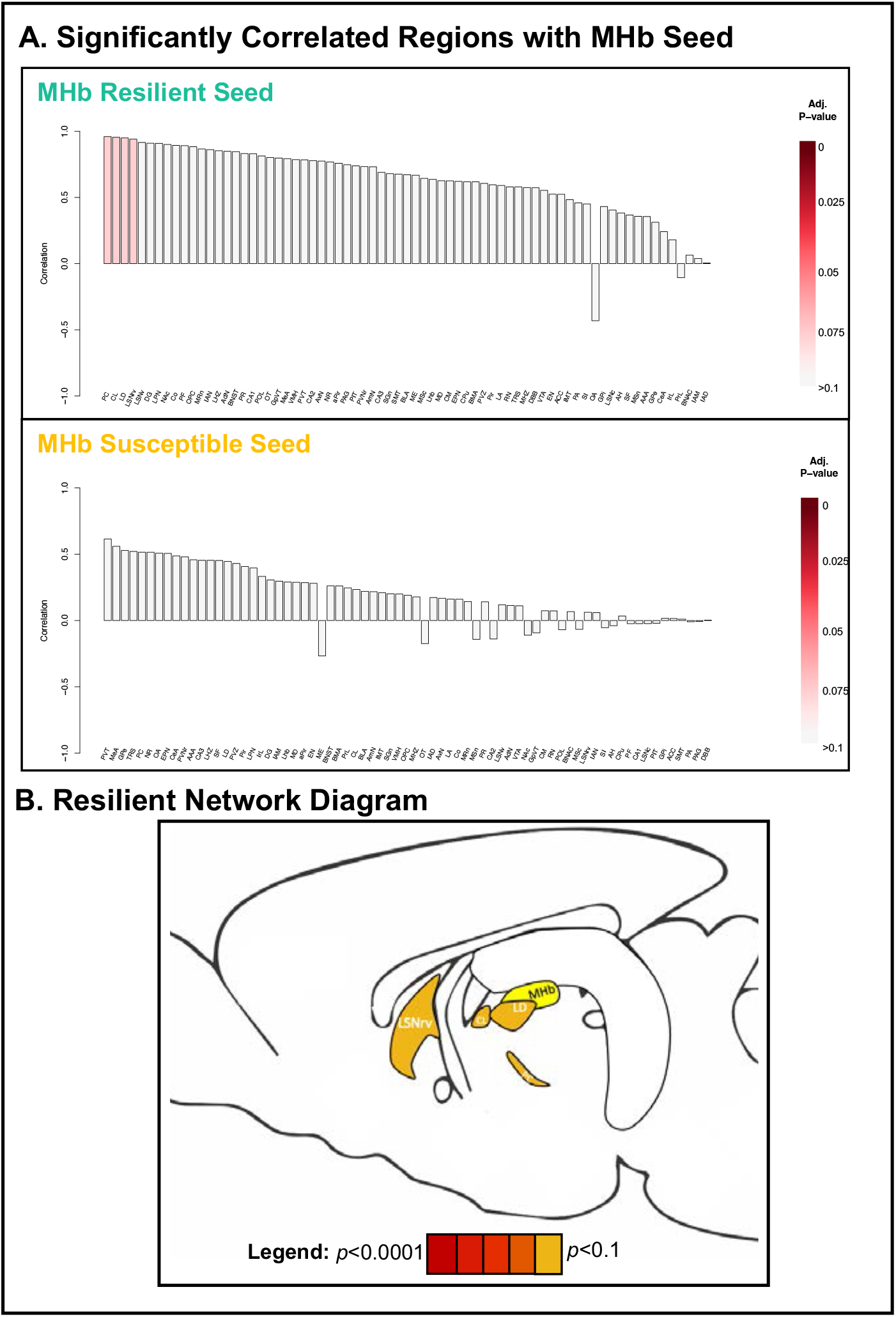
Significantly Correlated Brain Regions with MHb Seed. A. MHb Pearson R correlations of brain regions ranked based on adjusted p-values in future Resilient and Susceptible mice. B. Correlation network visualization of the significantly correlated regions with the MHb seed within future Resilient mice

#### Periventricular Zone Activity Correlates with Future SI Ratios

##### PVZ FosTRAP2+ Cells

While mice can be categorized as Resilient or Susceptible based on an SI Ratio of <1 or >1, we next wanted to ask whether the level of activation in a single brain region was correlated to the continuum of avoidance that an animal exhibited. We found that activation levels within the periventricular zone were positively correlated with Social Interaction Ratios, such that an increase in Day 1 activity in the periventricular zone was associated with increased Social Interaction on Day 11 (r=0.273, p=0.007) **(Fig 9)**. Of note, the PVZ was nominally significant in both our exploratory and candidate approach in our group comparisons of Resilient and Susceptible animals (exploratory: *p*=0.019, *FDR=*0.349, candidate: *p*=0.028, *FDR*=0.707). Therefore, analysis of FosTRAP2+ cells on the continuum of the social interaction ratio may provide an additional nuance in predicting the final social outcome.

**Figure 9.**
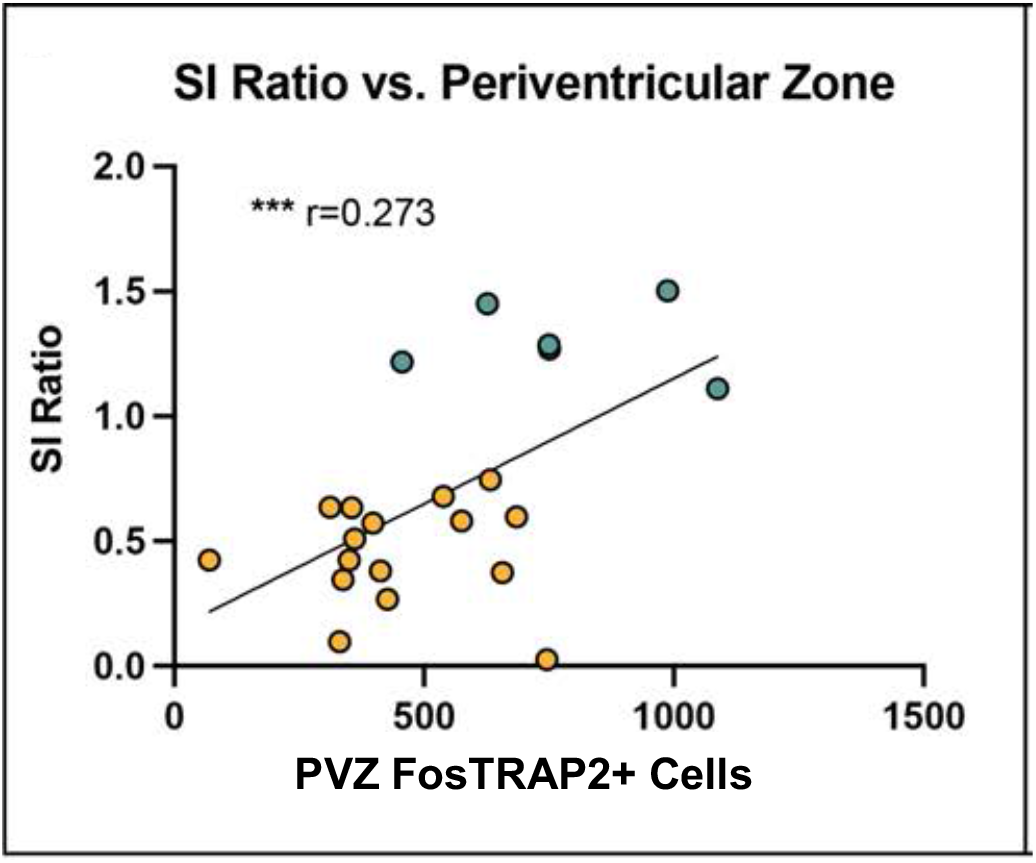
Correlates of PVZ FosTRAP2+ Brain Activity and Social Reactivity. A. SI Ratio has a significant positive correlation with number of FosTRAP+ counts within the Periventricular Zone (PVZ) (*r* = 0.273, *p*=0.007)

## DISCUSSION

Understanding how the brain responds to an initial defeat encounter may prove useful in elucidating how vulnerability to social stress arises. At baseline, before stress occurs, studies have assessed how microcircuits or specific brain regions selected *a priori* predict future states of vulnerability (Hultman et al. 2018; Muir et al. 2018). Many studies have focused on neural circuitry or activity patterns that elicit the social avoidance behavior after all 10 days of social defeat (Bagot et al. 2015; Chaudhury et al. 2013; Laine et al. 2017; Muir et al. 2020). This collective work demonstrates that the individual variation in the stress response manifests not only behaviorally in social avoidance, but likely as a coordinated neural response that engages multiple brain regions and networks of activation. Despite this, a key gap in our understanding of these neural responses has been the ability to gain resolution on unbiased brain-wide neural activation patterns that manifest during the first stress experience to predict future social outcome.

In our study, we used the transgenic Fos-TRAP2 system to ask whether brain-wide neural activation patterns during the initial defeat encounter predicts subsequent social-behavioral outcome. Overall, our findings demonstrate that the initial experience with social defeat induces a highly orchestrated brain-wide pattern of neural activation centering on key anatomical nodes that were revealed by two major approaches-within group and between group comparisons. Foremost among these regions are the lateral septum and the BNST that encoded defeat, the basomedial amygdala that encoded future susceptibility, and the hippocampal CA1 area and medial habenula that encoded future resilience. Moreover, activation in the PVZ on Day 1 correlated positively with future social interaction ratios.

In our assessment of 73 brain regions (exploratory) or 26 brain regions (candidate), the majority of brain areas exhibited similar levels of activation across Control and Defeated, as well as future Resilient and Susceptible groups. This could be due to a number of factors, as the initial experience of being placed in a novel social environment or in the Defeat encounter activates arousal, stress, as well as social systems (Martinez et al. 2002; Perkins et al. 2017; Tanimizu et al. 2017; VanElzakker et al. 2008). Only a small number of brain regions emerged as significantly different between groups.

However, when examining patterns of activation exhibited by each of the groups, as depicted by the heatmaps (**Fig. 4)**, it is evident that social defeat triggers a high degree of coordination across multiple brain regions relative to the control condition. In other words, animals who exhibit a strong response to defeat in one area are highly likely to exhibit a strong response in many other regions in a highly orchestrated manner. Equally remarkable is that the signature of social defeat is visibly different in the animals that will emerge as resilient vs. susceptible, with the resilient animals showing a more coherent response.

### The Bed Nucleus of the Stria Terminalis Activation Represents Features of Stress Modulation, Social Interaction, and Anxiety-Related Behaviors

The BNST is a region of high interest in psychiatric disorders, as it is known to be a sexually-dimorphic region that regulates a number of social and memory-related behaviors (Flanigan and Kash 2022). For example, regulation of fear behavior, social attachment behaviors, aggression-related behaviors, initiation of mating, and arousal have all been shown to be mediated through this highly anatomically connected region (Lebow and Chen 2016). In our Defeated animals, it appears that activation of the BNST most significantly correlates with several brain regions to which it has direct anatomical connections. Here, we will focus on a select number of regions that a) were significantly correlated with BNST activation; b) are known to have direct or indirect connections with the BNST; and c) are implicated in stress, social, and aggression circuits.

The BNST is a highly complex structure that, among other functions, plays a critical role in the central glucocorticoid negative feedback circuit that tracks and limits the stress response., The termination of the stress response arises in the hippocampus, with a glutamatergic projection to the BNST. In turn, BNST GABAergic neurons project directly to the PVN to inhibit the activity of corticotropin releasing factor (CRF) neurons and limit the stress response (Herman et al. 1995; Cullinan et al. 1993). Optogenetic stimulation of GABAergic BNST projections to parvalbumin nucleus accumbens neurons increases social interaction behavior and reduces anxiety-like behavior (Xiao et al. 2021).

In response to a social defeat, all 4 sub-regions of the hippocampus (CA1, CA2, CA3, and DG), as well as the PVN are highly correlated with BNST activity. Moreover, in Defeat animals, the BNST is significantly correlated with multiple amygdala regions. The BMA, basolateral amygdala (BLA), central amygdala (CeA), and medial amygdala (MeA) have all been shown to receive direct input from the BNST(Jasnow et al. 2004; Markham et al. 2009; Nordman et al. 2020; Walker and Davis 1997; Dong et al. 2001b). CRF projections from the BNST to the CeA go on to activate the HPA-axis, which is also modulated upstream through a projection from the BLA to the CeA. The BLA-CeA projection is known to be involved in processing threat-related information (Dong et al. 2001a; LeDoux 2007). The addition of the hippocampus and amygdala regions in the Defeat condition may represent a sustained stress response that is elicited through the physical and psychosocial stress component.

Nordman et al., found that the act of winning an agonistic encounter increased synaptic transmission between the MeA to the BNST and ventromedial hypothalamus (VMH) prior to subsequent encounters (Nordman et al. 2020). Indeed, in Defeat animals, the MeA and VMH are highly correlated with the BNST. The initial experience of Defeat therefore may be activating BNST-related circuits that involve the stress response, social cues, and threat-related assessments.

### Lateral Septal Nucleus, Ventral Part is Involved in Aggression-Related Behaviors

In addition to the BNST, the LSNv selectively increased activation in Defeat animals in comparison to Controls. Similar to the BNST, LS is a neurochemically diverse structure and acts as a relay station between multiple brain regions, including the prefrontal cortex, hippocampus, limbic regions (BLA, MeA, ventral tegmental area (VTA), BNST, and PVN) and transmits information to hypothalamic and thalamic regions to regulate an individual’s internal emotional and social state(Menon et al. 2022). As it is a hub between multiple brain regions, it is not surprising that within the Defeat group, LSNv activity is highly significantly correlated with several other areas.

The classic view of the lateral septum is that it inhibits aggression. Indeed, observations of “septal rage” septal tumors in humans were replicated in a wide range of animal studies (see Menon et al., 2022). Our findings show that LSNv activity is significantly correlated with activity in the VMH and CA2. This is of particular interest, as optogenetic activation of a projection from the CA2 to the LS has been used to disinhibit the VMH and trigger the termination of ongoing attacks during agonistic encounters (Lee et al. 2014; Wong et al. 2016). During the agonistic encounter, mice continuously engage and terminate their physical bouts of fight behavior. Therefore, this circuit may be modulating aspects of the aggression behavior in Defeat animals on Day 1. In addition to the CA2, LSNv activity is significantly correlated with activity in the CA1, CA3, and DG. In a resident-intruder paradigm in rats, an elevation of Fos protein expression was found in all 4 subregions of the dorsal and ventral hippocampus (Calfa et al. 2007). When a glucocorticoid antagonist was delivered into the LS prior to the final stress experience, the number of Fos+ cells throughout the hippocampus was reduced, in addition to a reduction in social avoidance behavior. This suggests that the LS mediates social avoidance via a glucocorticoid-sensitive mechanism (Calfa et al. 2007). It is therefore notable that in both the Control and Defeat conditions, the LS was the most highly significantly correlated region, as the modulation of social and emotional states appears to be highly dependent on the LS network.

### Between Group Analyses: Medial Habenula is Selectively Activated in Future Resilient Mice

In recent years, the habenula has been implicated in a number of psychiatric disorders, such as depression, ADHD, and schizophrenia (Hikosaka 2010; Lee and Goto 2011). Unlike the lateral habenula, which has been implicated in mediating reward and depression-like behavior, the role of the medial habenula (MHb) in these behaviors is less known. Lesions and optogenetic studies in the MHb have implicated it in exercise motivation, hedonic state, and reinforcement learning (Hsu et al. 2016, 2014).

Within the Resilient animals, a small network appeared to be correlated with the MHb. The posterior complex of the thalamus, a region involved in tactile-sensory integration was the most correlated, indicating that there may be some social-sensory related circuit relayed or activated in coordination with the MHb (Burton and Jones 1976; Casas-Torremocha et al. 2017). This network also comprises the centrolateral nucleus (CL) of the thalamus, which has a known connection with the MHb, and has been associated with attentional functions, as lesions to the CL cause metabolic cortical depression(Raos et al. 1995). Moreover, the lateraldorsal nucleus of the thalamus (LD), a relay region that provides inputs to higher order limbic-cortical areas was highly correlated with the MHb seed (Bezdudnaya and Keller 2008). In addition to the sensory-related regions, the LSNv, a region described above as being involved in social aggression, was also highly correlated with the MHb. This may seem contradictory at first, as activation of the LS is known to decrease aggression. However, with the temporal window of 4-OHT injection, we may be capturing features of the psychosocial stress component, in which the aggressive encounter is ended, and mice are placed across from the CD1 with no physical interaction possible. Therefore, we may be capturing future Resilient-specific circuits that integrate sensory-related information about the initial aggressive encounter.

### Distinct Network Hubs Relate to Future Resilience or Susceptibility

Our correlation network analyses identified distinct hubs in Future Resilient and Susceptible mice. In future Resilient mice, the hippocampal CA1 region emerged as a major node that significantly correlated 22 brain regions that passed FDR-correction. As part of the HPA axis, the CA1 sends GABAergic afferents to the BNST and PVN to mediate the stress response (Herman et al. 2005, 1995). Moreover, activity within the CA1 region has been shown to be differentially attuned to acute versus chronic stressors. In a longitudinal study where local field potentials were recorded within the CA1 region on the first and last day of chronic stress, researchers found that the initial stress experience increased firing in the CA1, while repeated exposure to stress led to decreased firing (Tomar et al. 2021). Without the temporal resolution, we do not have clarity on the specific activation pattern of the CA1, however, it is clearly highly involved in the initial acute stress response of future Resilient mice.

In future Susceptible mice, the BMA emerged as a major node that significantly correlated with 52 brain regions that passed FDR correction. Electrophysiology and optogenetics studies have demonstrated that selective populations within the BMA code aversive or safe environments (Sangha et al. 2013). In addition, recordings within the BMA during controllable and uncontrollable stress revealed that uncontrollable stress leads to higher BMA activity than a controllable stress stimulus (Adhikari et al. 2015). Auditory and visual information about predators is also processed through the BMA, as lesions to the BMA have been shown to reduce freezing behaviors in response to predator pheromone exposure (decreased defensive coping) (Martinez et al. 2011). Therefore, the node of activation within the BMA in future Susceptible mice is possibly associated with threat and anxiety-like states that may set the course for future vulnerability.

### Greater Activation of the Periventricular Zone (PVZ) is Associated with Future Resilience

We found that individuals who exhibited a higher level of social interaction following stress, and subsequently greater resilience, had a higher number of FosTRAP+ cell counts within the PVZ on Day1. In our previous work (Murra et al. 2022), we found that there was a significant correlation between corticosterone levels and increased social interaction when mice were placed back in a social stress context (the forced interaction test) following social defeat. We hypothesize that increased PVZ activity during the first social defeat encounter reflects a robust hormonal stress response that was maintained after repeated defeat. The PVZ consists of both parvocellular and magnocellular neurosecretory cells. Parvocellular cells express corticotropin releasing hormone (CRH), while magnocellular cells express oxytocin and vasopressin (Dudas 2013). Future studies that explore which cell type co-localizes with these PVZ FosTRAP+ cells may reveal how initial activation of these stress or social-related cell-types mediate the final social outcome.

### Conclusion

Overall, the FosTRAP2 approach has captured a rich network of activation that appears to include sensory, emotional, motivational, and compensatory mechanisms of coping with a social encounter. For the first time, our study has identified brain regions that are selective to the initial experience of Defeat, as well as Resilient or Susceptible outcomes. In addition, we have identified networks of brain regions that correlated highly with seed regions we have identified and revealed brain-wide activation patterns. Therefore, we can conclude that there are brain-wide neural signatures that are present during the initial defeat encounter that map onto the stress experience. Further, that these signatures are distinct from the social-only (control) experience and may reflect future Susceptible and Resilient outcomes. Our work provides an important framework for future studies to explore the biochemical identity and circuit mechanisms behind key brain regions and their ability to mediate resilient or susceptible outcomes.

## ACKNOWLEDGMENTS

This work was supported by the following grants: Hope for Depression Research Foundation (HDRF); Office of Naval Research (ONR) N00014-19-1-2149; The Pritzker Neuropsychiatric Research Consortium. We would also like to acknowledge Vivek Kumar, David Krolewski, and Matthew Foltz for discussions and guidance on imaging and image analysis. In addition, Megan Hagenauer for consultation on statistical considerations and Claudia Laborc, Qiang Wei, and Limei Zhang for mouse colony management.

