## Supplement for "Patterns of Neural Activation During an Initial Social Stress Encounter are Predictive of Future Susceptibility or Resilience: A FosTRAP2 Study"

**SUPPLEMENTAL**

*Table S1: Brain Regions Selected for Exploratory Approach (73)*

| Acronym | Name |
|---|---|
| ACC | Anterior cingulate area |
| PL | Prelimbic area |
| IL | Infralimbic area |
| OA | Orbital area |
| Pir | Piriform area |
| Co | Cortical amygdalar area |
| aPir | Piriform-amygdalar area |
| CA1 | Field CA1 |
| CA2 | Field CA2 |
| CA3 | Field CA3 |
| DG | Dentate gyrus |
| EPN | Endopiriform nucleus |
| LA | Lateral amygdalar nucleus |
| BLA | Basolateral amygdalar nucleus |
| BMA | Basomedial amygdalar nucleus |
| PA | Posterior amygdalar nucleus |
| CPu | Caudoputamen |
| NAc | Nucleus accumbens |
| OT | Olfactory tubercle |
| LSNc | Lateral septal nucleus/ caudal (caudodorsal) part |
| LSNrv | Lateral septal nucleus/ rostral (rostroventral) part |
| LSNv | Lateral septal nucleus/ ventral part |
| SF | Septofimbrial nucleus |
| AAA | Anterior amygdalar area |
| BNAOt | Bed nucleus of the accessory olfactory tract |
| CeA | Central amygdalar nucleus |
| IAN | Intercalated amygdalar nucleus |
| MeA | Medial amygdalar nucleus |
| GPe | Globus pallidus/ external segment |
| GPi | Globus pallidus/ internal segment |
| SI | Substantia innominata |
| MRn | Magnocellular nucleus |
| MSc | Medial septal complex |
| MSn | Medial septal nucleus |
| DBB | Diagonal band nucleus |
| TRS | Triangular nucleus of septum |
| BNST | Bed nuclei of the stria terminalis |
| BNAC | Bed nucleus of the anterior commissure |
| LPN | Lateral posterior nucleus of the thalamus |
| PC | Posterior complex of the thalamus |
| POL | Posterior limiting nucleus of the thalamus |
| SGn | Suprageniculate nucleus |
| EN | Ethmoid nucleus of the thalamus |
| AvN | Anteroventral nucleus of thalamus |
| AmN | Anteromedial nucleus |
| AdN | Anterodorsal nucleus |

| | |
|---|---|
| IAM | Interanteromedial nucleus of the thalamus |
| IAD | Interanterodorsal nucleus of the thalamus |
| LD | Lateral dorsal nucleus of thalamus |
| IMT | Intermediodorsal nucleus of the thalamus |
| MD | Mediodorsal nucleus of thalamus |
| SMT | Submedial nucleus of the thalamus |
| PR | Perireunensis nucleus |
| PVT | Paraventricular nucleus of the thalamus |
| NR | Nucleus of reuniens |
| RN | Rhomboid nucleus |
| CM | Central medial nucleus of the thalamus |
| OPC | Paracentral nucleus |
| CL | Central lateral nucleus of the thalamus |
| PF | Parafascicular nucleus |
| PIT | Posterior intralaminar thalamic nucleus |
| GpVT | Geniculate group/ ventral thalamus |
| Mhb | Medial habenula |
| Lhb | Lateral habenula |
| PVZ | Periventricular zone |
| PVNr | Periventricular region |
| MHZ | Hypothalamic medial zone |
| LHZ | Hypothalamic lateral zone |
| VMH | Ventromedial hypothalamic nucleus |
| AH | Anterior hypothalamic nucleus |
| ME | Median eminence |
| VTA | Ventral tegmental area |
| PAG | Periaqueductal gray |

*Table S2: Brain Regions Selected for Candidate Approach (26)*

| **Acronym** | **Brain Region** |
|---|---|
| PL | Prelimbic area |
| IL | Infralimbic area |
| CA1 | Field CA1 |
| CA2 | Field CA2 |
| CA3 | Field CA3 |
| DG | Dentate gyrus |
| LA | Lateral amygdalar nucleus |
| BMA | Basomedial amygdalar nucleus |
| Cpu | Caudoputamen |
| LSNc | Lateral septal nucleus/ caudal (caudodorsal) part |
| LSNr | Lateral septal nucleus/ rostral (rostroventral) part |
| LSNv | Lateral septal nucleus/ ventral part |
| CeA | Central amygdalar nucleus |
| IAN | Intercalated amygdalar nucleus |
| MeA | Medial amygdalar nucleus |
| BNST | Bed nuclei of the stria terminalis |
| PVT | Paraventricular nucleus of the thalamus |
| MHb | Medial habenula |

| | |
|---|---|
| LHb | Lateral habenula |
| PVH | Periventricular Region |
| VMH | Ventromedial hypothalamic nucleus |
| NAc | Nucleus accumbens |
| VTA | Ventral tegmental area |
| LHZ | Hypothalamic lateral zone |
| GVT | Geniculate group/ ventral thalamus |
| BLA | Basolateral amygdalar nucleus |

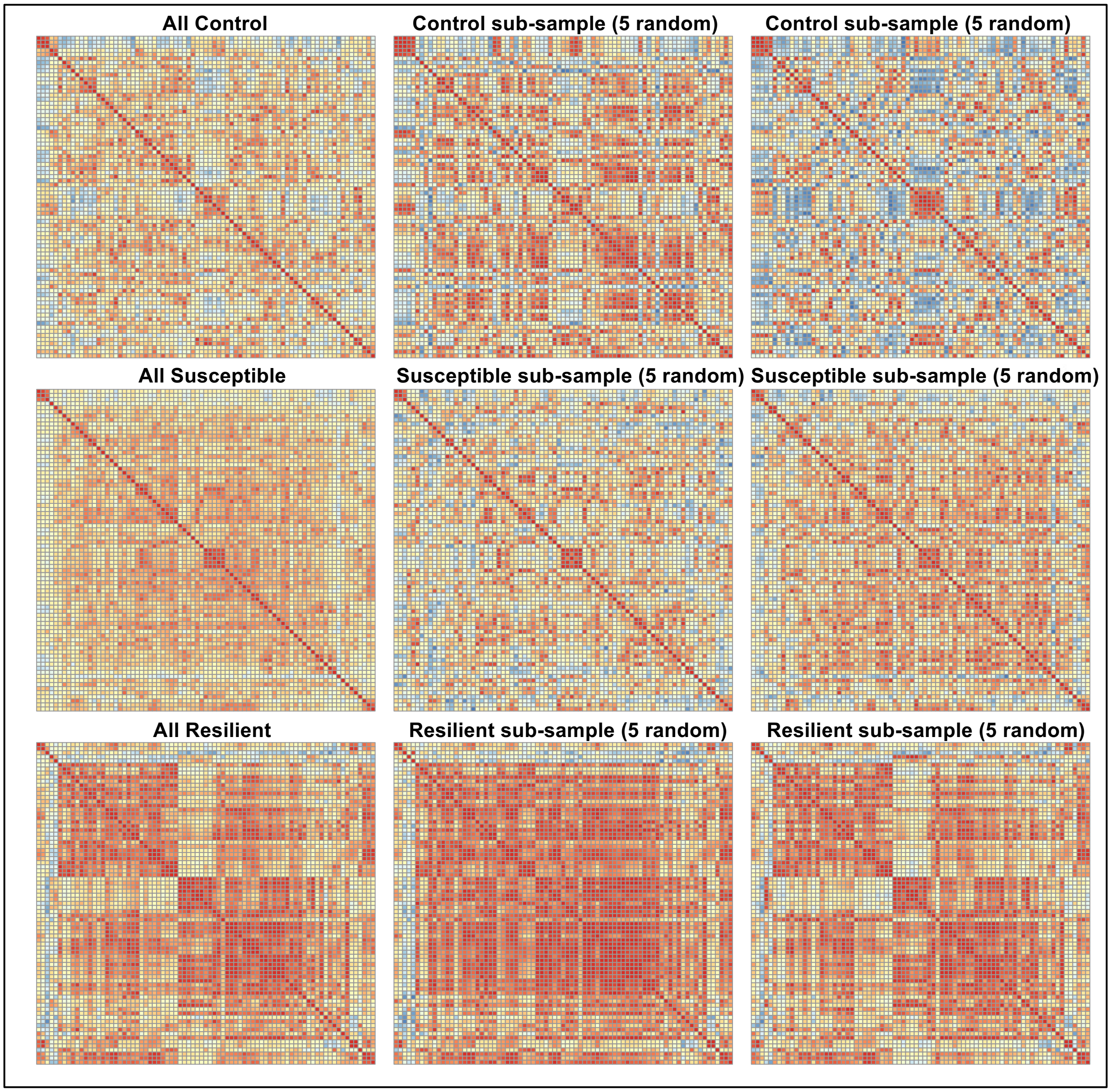


*Figure S1 Heatmap of Sub-Sampling 5 Random Animals Per Group*
Visual patterns are similar to what is observed when all animals are included, indicating that the low sample size within

the Resilient animals represents a real pattern of increased activation compared to Susceptible and Control groups. Colors in heatmap indicate a range of Pearson *r* correlations, more red=1, more blue=-1.

**Defeat vs. Control BNST Seed:**

*Table S3 Top Brain Regions Correlated with the BNST in Defeated Mice*

| Region | Correlation | P.Value | FDR |
|---|---|---|---|
| CPu | 0.7823 | 0.0000 | 0.0012 |
| PVNr | 0.7576 | 0.0000 | 0.0016 |
| MRn | 0.7018 | 0.0003 | 0.0047 |
| ACC | 0.6984 | 0.0003 | 0.0047 |
| EPN | 0.6954 | 0.0003 | 0.0047 |
| MHZ | 0.6736 | 0.0006 | 0.0071 |
| BMA | 0.6575 | 0.0009 | 0.0082 |
| NAc | 0.6552 | 0.0009 | 0.0082 |
| AvN | 0.6514 | 0.001 | 0.0082 |
| LSNrv | 0.6418 | 0.0013 | 0.0091 |
| Pir | 0.6384 | 0.0014 | 0.0091 |
| LHZ | 0.6268 | 0.0018 | 0.0106 |
| LSNv | 0.6214 | 0.002 | 0.0106 |
| SI | 0.6206 | 0.0021 | 0.0106 |
| CA2 | 0.6146 | 0.0023 | 0.0112 |
| IAN | 0.593 | 0.0036 | 0.0156 |
| MSc | 0.5909 | 0.0038 | 0.0156 |
| OT | 0.5894 | 0.0039 | 0.0156 |
| GPe | 0.5856 | 0.0042 | 0.0156 |
| PVZ | 0.5839 | 0.0043 | 0.0156 |
| DBB | 0.5808 | 0.0046 | 0.0157 |
| AmN | 0.575 | 0.0051 | 0.0167 |
| PC | 0.5625 | 0.0064 | 0.0185 |
| LPN | 0.5621 | 0.0065 | 0.0185 |
| aPir | 0.5617 | 0.0065 | 0.0185 |
| SF | 0.5578 | 0.007 | 0.0185 |
| PVT | 0.5561 | 0.0072 | 0.0185 |
| AdN | 0.556 | 0.0072 | 0.0185 |
| MeA | 0.5418 | 0.0092 | 0.0228 |
| CM | 0.5377 | 0.0099 | 0.0234 |
| POL | 0.5364 | 0.0101 | 0.0234 |
| CeA | 0.5285 | 0.0115 | 0.0258 |
| CA1 | 0.5251 | 0.0121 | 0.0264 |
| Co | 0.5205 | 0.013 | 0.0275 |
| DG | 0.5184 | 0.0135 | 0.0277 |
| BLA | 0.5005 | 0.0177 | 0.0354 |
| GpVT | 0.491 | 0.0203 | 0.0395 |
| AH | 0.4844 | 0.0223 | 0.0423 |
| VMH | 0.465 | 0.0292 | 0.0518 |

| | | | |
|---|---|---|---|
| AAA | 0.4647 | 0.0293 | 0.0518 |
| IAD | 0.4644 | 0.0295 | 0.0518 |
| VTA | 0.4609 | 0.0309 | 0.0529 |
| CA3 | 0.4446 | 0.0382 | 0.0639 |
| OPC | 0.4386 | 0.0412 | 0.0674 |
| LD | 0.4284 | 0.0467 | 0.0724 |
| Lhb | 0.4278 | 0.047 | 0.0724 |
| LSNc | 0.4274 | 0.0473 | 0.0724 |
| TRS | 0.419 | 0.0523 | 0.0784 |
| Mhb | 0.4138 | 0.0556 | 0.0816 |
| PAG | 0.4076 | 0.0597 | 0.0859 |
| MD | 0.405 | 0.0615 | 0.0869 |
| PA | 0.3978 | 0.0667 | 0.0924 |
| PF | 0.3887 | 0.0738 | 0.1003 |

Red meets FDR cut-off, Brown is nominally significant. Black is non-significant

*Table S4 Top Brain Regions Correlated with the BNST in Control Mice*

| Region | Correlation | P.Value | FDR |
|---|---|---|---|
| PVNr | 0.8678 | 0.0024 | 0.1746 |
| ACC | 0.8090 | 0.0083 | 0.2383 |
| SI | 0.6760 | 0.0456 | 0.6098 |
| NAc | 0.6703 | 0.0487 | 0.6098 |
| PIT | -0.6421 | 0.0622 | 0.6098 |
| MSn | 0.6161 | 0.0773 | 0.6098 |
| AAA | 0.6131 | 0.0792 | 0.6098 |
| Pir | 0.5717 | 0.1078 | 0.6098 |

Red meets FDR cut-off, Brown is nominally significant. Black is non-significant

**Defeat vs. Control LSNv Seed:**

*Table S5 Top Brain Regions Correlated with the LSNv in Defeated Mice*

| Region | Correlation | P.Value | FDR |
|---|---|---|---|
| LSNrv | 0.9153 | 0.0000 | 0.0000 |
| NAc | 0.8105 | 0.0000 | 0.0002 |
| OT | 0.7902 | 0.0000 | 0.0003 |
| ACC | 0.738 | 0.0000 | 0.0014 |
| LPN | 0.7346 | 0.0000 | 0.0014 |
| VMH | 0.7112 | 0.0002 | 0.0021 |
| Pir | 0.7108 | 0.0002 | 0.0021 |
| PC | 0.6986 | 0.0003 | 0.0027 |
| DG | 0.6914 | 0.0004 | 0.0029 |
| LHZ | 0.6877 | 0.0004 | 0.0029 |
| SI | 0.6831 | 0.0005 | 0.003 |
| MSc | 0.6741 | 0.0006 | 0.0035 |
| NR | 0.6532 | 0.0010 | 0.0054 |

| | | | |
|---|---|---|---|
| MeA | 0.6485 | 0.0011 | 0.0056 |
| CPu | 0.6428 | 0.0013 | 0.006 |
| CL | 0.6252 | 0.0019 | 0.0084 |
| BNST | 0.6214 | 0.0020 | 0.0086 |
| CA1 | 0.6167 | 0.0022 | 0.0089 |
| PF | 0.5978 | 0.0033 | 0.0114 |
| MSn | 0.5976 | 0.0033 | 0.0114 |
| RN | 0.5951 | 0.0035 | 0.0114 |
| PVNr | 0.593 | 0.0036 | 0.0114 |
| BMA | 0.5911 | 0.0038 | 0.0114 |
| GpVT | 0.5888 | 0.0039 | 0.0114 |
| EPN | 0.5868 | 0.0041 | 0.0114 |
| AvN | 0.5866 | 0.0041 | 0.0114 |
| PR | 0.5733 | 0.0053 | 0.0141 |
| PAG | 0.5679 | 0.0058 | 0.015 |
| CM | 0.564 | 0.0063 | 0.0154 |
| LD | 0.5625 | 0.0064 | 0.0154 |
| IMT | 0.5545 | 0.0074 | 0.0172 |
| MD | 0.5497 | 0.008 | 0.0181 |
| MHZ | 0.5473 | 0.0084 | 0.0183 |
| BLA | 0.5436 | 0.0089 | 0.0189 |
| CA2 | 0.5399 | 0.0095 | 0.0192 |
| DBB | 0.5394 | 0.0096 | 0.0192 |
| POL | 0.5349 | 0.0103 | 0.0201 |
| AmN | 0.5229 | 0.0125 | 0.0237 |
| CA3 | 0.5195 | 0.0132 | 0.0244 |
| PVT | 0.5139 | 0.0144 | 0.026 |
| Co | 0.4909 | 0.0204 | 0.0358 |
| MRn | 0.4766 | 0.0249 | 0.0428 |
| OPC | 0.4705 | 0.0271 | 0.0451 |
| LA | 0.4689 | 0.0277 | 0.0451 |
| SF | 0.4677 | 0.0282 | 0.0451 |
| EN | 0.4408 | 0.0401 | 0.0627 |
| AH | 0.4371 | 0.0419 | 0.0642 |
| AdN | 0.4156 | 0.0544 | 0.0816 |
| PVZ | 0.4118 | 0.0569 | 0.0828 |
| Mhb | 0.4109 | 0.0575 | 0.0828 |
| SMT | 0.4037 | 0.0624 | 0.0881 |
| PA | 0.3887 | 0.0738 | 0.1022 |
| VTA | 0.3595 | 0.1004 | 0.1363 |

Red meets FDR cut-off, Brown is nominally significant, and Black is non-significant

*Table S6 Top Brain Regions Correlated with the LSNv in Control Mice*

| Region | Correlation | P.Value | FDR |
|---|---|---|---|
| AmN | 0.8906 | 0.0013 | 0.0912 |
| LSNrv | 0.8534 | 0.0034 | 0.1223 |
| OPC | 0.8350 | 0.0051 | 0.1223 |
| AH | 0.8153 | 0.0074 | 0.1334 |
| EPN | 0.7939 | 0.0106 | 0.1529 |
| RN | 0.7638 | 0.0166 | 0.1614 |
| CPu | 0.7599 | 0.0175 | 0.1614 |
| LHZ | 0.7581 | 0.0179 | 0.1614 |
| CeA | 0.7371 | 0.0235 | 0.1717 |
| AAA | 0.7202 | 0.0287 | 0.1717 |
| MHZ | 0.719 | 0.029 | 0.1717 |
| AvN | 0.7155 | 0.0302 | 0.1717 |
| SF | 0.7132 | 0.031 | 0.1717 |
| ACC | 0.6979 | 0.0366 | 0.1736 |
| CM | 0.6934 | 0.0383 | 0.1736 |
| AdN | 0.6928 | 0.0386 | 0.1736 |
| PF | 0.6811 | 0.0434 | 0.1780 |
| PVZ | 0.6786 | 0.0445 | 0.1780 |
| DG | 0.6627 | 0.0517 | 0.1859 |

Red meets FDR cut-off, Brown is nominally significant, and Black is non-significant

**MHb Seed:**

*Table S7 Top Brain Regions Correlated with the MHb in Future Resilient*

| Region | Correlation | P.Value | FDR |
|---|---|---|---|
| PC | 0.9597 | 0.0024 | 0.0997 |
| CL | 0.9544 | 0.0031 | 0.0997 |
| LD | 0.9497 | 0.0037 | 0.0997 |
| LSNrv | 0.9402 | 0.0053 | 0.0997 |
| LSNv | 0.9154 | 0.0104 | 0.1180 |
| DG | 0.9096 | 0.0119 | 0.1180 |
| LPN | 0.9085 | 0.0122 | 0.1180 |
| NAc | 0.9002 | 0.0145 | 0.1180 |
| Co | 0.8934 | 0.0165 | 0.1180 |
| PF | 0.8911 | 0.0171 | 0.1180 |
| OPC | 0.8832 | 0.0197 | 0.1180 |
| MRn | 0.8663 | 0.0256 | 0.1418 |
| IAN | 0.8609 | 0.0277 | 0.1424 |
| LHZ | 0.8529 | 0.0309 | 0.1432 |
| AdN | 0.8484 | 0.0327 | 0.1432 |
| BNST | 0.8458 | 0.0338 | 0.1432 |
| PR | 0.8317 | 0.0401 | 0.1556 |
| CA1 | 0.8296 | 0.0411 | 0.1556 |
| POL | 0.8134 | 0.049 | 0.1763 |

| | | | |
|---|---|---|---|
| OT | 0.8018 | 0.055 | 0.1867 |

Red meets FDR cut-off, Brown is nominally significant, and Black is non-significant

*Table S8 Top Brain Regions Correlated with the MHb in Future Susceptible Mice*

| Region | Correlation | P.Value | FDR |
|---|---|---|---|
| PVT | 0.6146 | 0.0113 | 0.4023 |
| MeA | 0.5595 | 0.0242 | 0.4023 |
| GPe | 0.5284 | 0.0354 | 0.4023 |
| TRS | 0.5211 | 0.0385 | 0.4023 |
| PC | 0.5151 | 0.0411 | 0.4023 |
| NR | 0.5149 | 0.0413 | 0.4023 |
| OA | 0.5075 | 0.0448 | 0.4023 |
| EPN | 0.5063 | 0.0454 | 0.4023 |
| CeA | 0.4872 | 0.0556 | 0.4023 |
| PVNr | 0.4804 | 0.0596 | 0.4023 |

Brown is nominally significant, and Black is non-significant
